## Supplementary figures and images for "Therapeutic effects of PDGF-AB/BB against cellular senescence in human intervertebral disc"

### Supplemental Figure 1

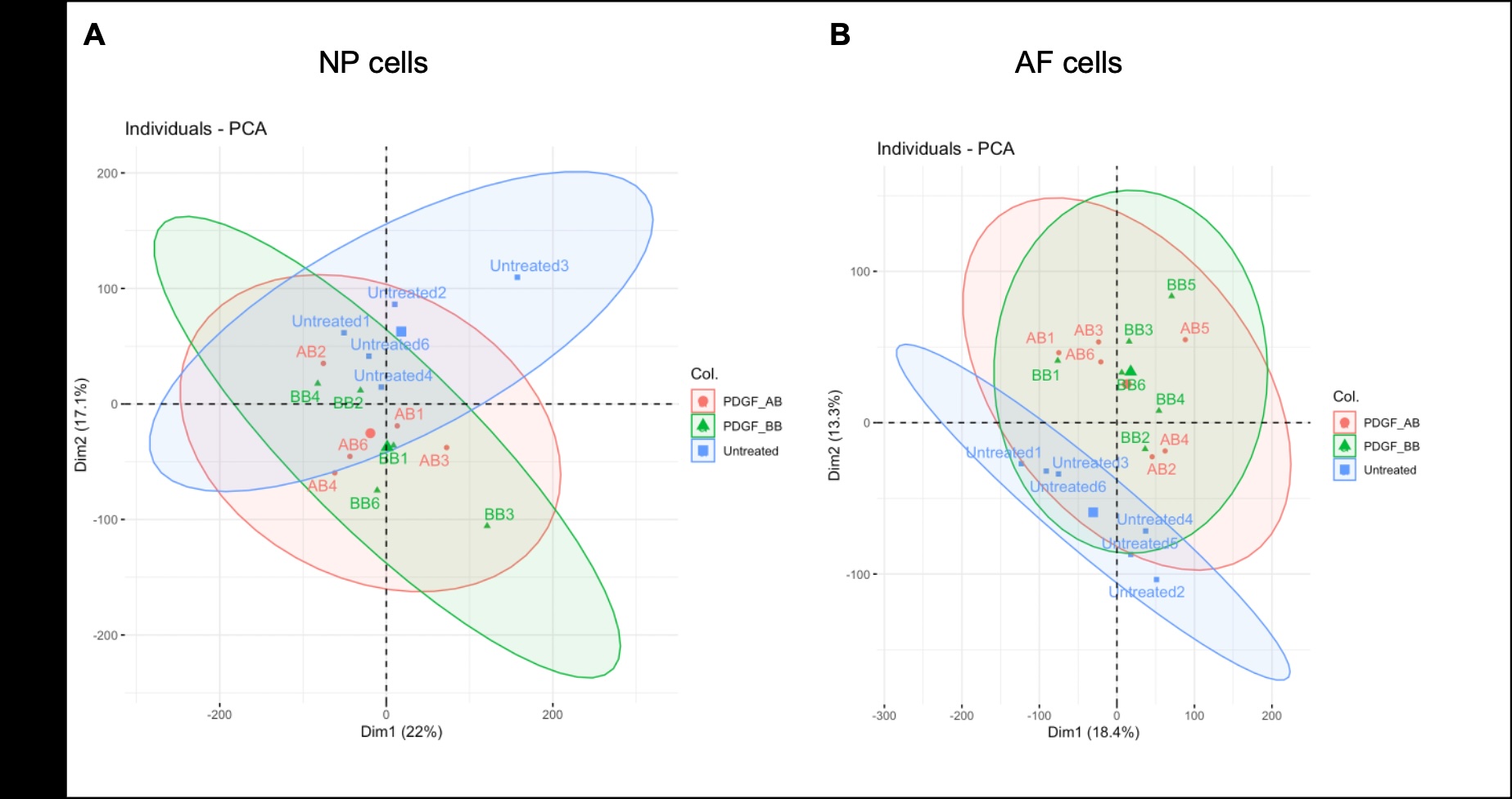

### Supplemental Figure 2

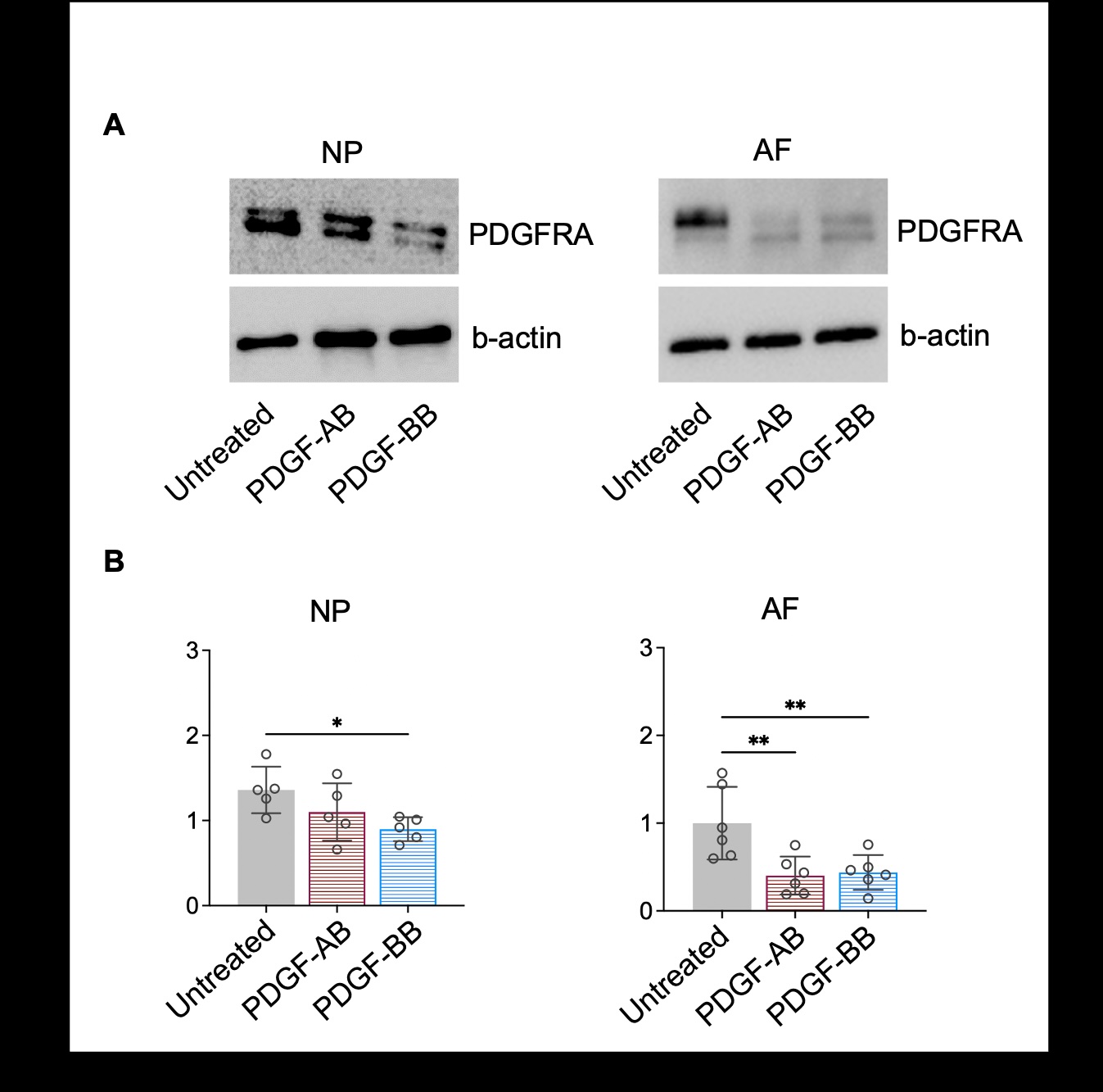
